# Inducible expression of (pp)pGpp synthetases in *Staphylococcus aureus* is associated with activation of stress response genes

**DOI:** 10.1101/2020.04.25.059725

**Authors:** Petra Horvatek, Andrew Magdy Fekry Hanna, Fabio Lino Gratani, Daniela Keinhörster, Natalya Korn, Marina Borisova, Christoph Mayer, Dominik Rejman, Ulrike Mäder, Christiane Wolz

**Affiliations:** Interfaculty Institute of Microbiology and Infection Medicine, University of Tuebingen, Germany; Quantitative Proteomics & Proteome Center Tuebingen, University of Tuebingen, Germany; Institute of Organic Chemistry and Biochemistry, Czech Academy of Sciences, Prague, Czech Republic; Interfaculty Institute for Genetics and Functional Genomics, University Medicine Greifswald, Greifswald

**Keywords:** Stringent response, (p)ppGpp, pppGpp, RSH, oxidative stress, PSM, *Staphylococcus aureus*

## Abstract

The stringent response is characterized by the synthesis of the messenger molecules pppGpp, ppGpp or pGpp (here collectively designated (pp)pGpp). The phenotypic consequences resulting from (pp)pGpp accumulation vary among species and can be mediated by different underlying mechanisms. Most genome-wide analyses have been performed under stress conditions, which often mask the immediate effects of (pp)pGpp-mediated regulatory circuits. In *Staphylococcus aureus*, (pp)pGpp can be synthesized via the RelA-SpoT-homolog (RSH_*Sau*_) upon amino acid limitation or via one of the two small (pp)pGpp synthetases RelP or RelQ, upon cell wall stress. We used RNA-Seq to compare the global effects in response to transcriptional induction of the synthetase domain of RSH (RSH-Syn), RelP or RelQ without the need to apply additional stress conditions. Enzyme expression resulted in changes in the nucleotide pool similar to induction of the stringent response via the tRNA synthetase inhibitor mupirocin: a reduction in the GTP pool, an increase in the ATP pool and synthesis of pppGpp, ppGpp and pGpp. Induction of all three enzymes resulted in similar changes in the transcriptome. However, RelQ was less active than RSH-Syn and RelP, indicating strong restriction of its (pp)pGpp-synthesis activity *in vivo*. Genes involved in the SOS response, iron storage (e.g. *ftnA, dps*), oxidative stress response (e.g., *katA, sodA*) and the the *psmα1-4 and psmß1-2* operons coding for cytotoxic, phenole soluble modulins (PSMs) were highly upregulated upon (pp)pGpp synthesis. Analyses of the *ftnA, dps* and *psm* genes in different regulatory mutants revealed that their (pp)pGpp-dependent regulation can occur independent of the regulators PerR, Fur, SarA or CodY. Moreover, *psm* expression is uncoupled from expression of the quorum sensing system Agr, the main known *psm* activator. The expression of central genes of the oxidative stress response protects the bacteria from anticipated ROS stress derived from PSMs or exogenous sources. Thus, we identified a new link between the stringent response and oxidative stress in *S. aureus* that is likely crucial for survival upon phagocytosis.

**Significance:** Most bacteria make use of the second messenger (pp)pGpp to reprogram bacterial metabolism under nutrient-limiting conditions. In the human pathogen *Staphylococcus aureus*, (pp)pGpp plays an important role in virulence, phagosomal escape and antibiotic tolerance. Here, we analyzed the immediate consequences of (pp)pGpp synthesis upon transcriptional induction of the (pp)pGpp-producing enzymes RSH, RelP or RelQ. (pp)pGpp synthesis provokes immediate changes in the nucleotide pool and severely impacts the expression of hundreds of genes. A main consequence of (pp)pGpp synthesis in *S. aureus* is the induction of ROS-inducing toxic phenol-soluble modulins (PSMs) and simultaneous expression of the detoxifying system to protect the producer. This mechanism is likely of special advantage for the pathogen after phagocytosis.

## Introduction

The stringent response is characterized by the synthesis of the alarmones pGpp, ppGpp and pppGpp, here collectively named (pp)pGpp. (pp)pGpp interferes with many cellular processes, including transcription, replication and translation (Irving & Corrigan, 2018, Dozot *et al*., 2006, Steinchen & Bange, 2016, Wu & Xie, 2009, Zhu *et al*., 2019, Gaca *et al*., 2015a, Liu *et al*., 2015, Hobbs & Boraston, 2019, Wolz *et al*., 2010, Hauryliuk *et al*., 2015). Depending on the species, the stringent response is crucial for diverse biological processes, including differentiation, biofilm formation, antibiotic tolerance, production of secondary metabolites or virulence (Dalebroux *et al*., 2010, Hobbs & Boraston, 2019). It is now clear that there are fundamental differences between the stringent response initially characterized in *E. coli* and the stringent response in Firmicutes (Wolz *et al*., 2010, Liu *et al*., 2015). Differences have been observed in the enzymes involved in the synthesis and degradation of the messengers and in the downstream effects of (pp)pGpp.

(pp)pGpp is synthesized by long RelA-SpoT-homologs (RSHs) or small alarmone synthetases (SAS) by transferring pyrophosphate originating from ATP to the 3’ OH group of GTP, GDP or GMP. RSH enzymes are present in nearly all bacteria and show a conserved molecular architecture composed of a C-terminal sensory domain and an N-terminus with distinct (pp)pGpp hydrolase and synthetase domains (Atkinson *et al*., 2011). Firmicutes, such as *Staphylococcus aureus*, possess one bifunctional RSH enzyme and one or two SAS enzymes, RelP and RelQ. Of note, bifunctional RSH enzymes are also named Rel or RelA. Amino acid limitation is the only condition known to induce an RSH-mediated stringent response phenotype (Geiger *et al*., 2010). Under non-inducing conditions, RSH_*Sau*_ is primarily in a hydrolase-On/synthetase-Off conformation even when the C-terminal sensory domain is deleted (Gratani, Horvatek *et al*. 2018). The strong hydrolase activity of RSHs makes these enzymes essential for the detoxification of (pp)pGpp produced by RelP or RelQ (Geiger *et al*., 2010).

The small SAS enzymes in *S. aureus* are part of the VraRS cell-wall stress regulon (Kuroda *et al*., 2003) and are thus transcriptionally induced, e.g., after vancomycin treatment (Geiger *et al*., 2014). Thereby, they contribute to tolerance towards cell-wall active antibiotics such as ampicillin or vancomycin. Recently, structural and mechanistic characterization revealed that RelQ from *Bacillus subtilis* and *Enterococcus faecalis* form tetramers (Steinchen *et al*., 2015, Beljantseva *et al*., 2017). RelQ activity is strongly inhibited through the binding of single-stranded RNA. pppGpp binding leads to disassociation of the RelQ:RNA complex and its activation (Beljantseva *et al*., 2017). In contrast, RelP activity is inhibited by both pppGpp and ppGpp, activated by Zn^2+^ and insensitive to inhibition by RNA (Manav *et al*., 2018, Steinchen *et al*., 2018). Thus, although highly homologous, RelP and RelQ seem to have different functions within the cell. One can assume that different post-translational regulatory mechanisms are in play to fine-tune (pp)pGpp synthesis under different growth conditions.

In *S. aureus*, the stringent response plays important roles in virulence (Geiger *et al*., 2010), phagosomal escape (Geiger *et al*., 2012) and antibiotic tolerance (Geiger *et al*., 2014, Corrigan *et al*., 2016, Gao *et al*., 2010, Dordel *et al*., 2014, Matsuo *et al*., 2019, Katayama *et al*., 2017, Hobbs & Boraston, 2019). The enzymes HprT and Gmk involved in GTP synthesis, putative GTPases (RsgA, RbgA, Era, HflX, and ObgE) and DNA primase were identified as (pp)pGpp target proteins (Kriel *et al*., 2012, Corrigan *et al*., 2016)(Wang *et al*., 2007). (pp)pGpp binding results in inhibition of these proteins, resulting in lowering of the GTP pool and inhibition of the translation apparatus and replication. Of note, in contrast to *E. coli*, (pp)pGpp from Firmicutes does not interfere with RNA polymerase activity (Hauryliuk *et al*., 2015). Instead, in these organisms, (pp)pGpp regulates transcription via an indirect mechanism that strongly relies on the lowering of the intracellular GTP pool (Kriel *et al*., 2012, Geiger *et al*., 2012, Krasny *et al*., 2008). A decrease in the GTP level leads to the repression of nucleotide-sensitive, GTP-initiating promoters, e.g., those of rRNA genes (Kastle *et al*., 2015, Krasny & Gourse, 2004). Low GTP levels also affect the CodY regulon. The transcription factor CodY, when loaded with GTP and branched-chain amino acids, acts mainly as a repressor of many genes involved in amino acid synthesis and virulence (Pohl *et al*., 2009, Majerczyk *et al*., 2010). The global transcriptional effects of (pp)pGpp have been examined previously in several Firmicutes, such as *B. subtilis* (Eymann *et al*., 2002), *Streptococcus pneumoniae* (Kazmierczak *et al*., 2009), *Enterococcus faecalis* (Gaca *et al*., 2012), *Streptococcus mutans* (Nascimento *et al*., 2008) and *S. aureus* (Geiger *et al*., 2012). These studies were based on the comparison of the wild-type and RSH mutant strains under conditions mimicking amino acid starvation. Of note, these stress conditions are accompanied by profound physiological changes, which are only partially mediated by (pp)pGpp. For instance, amino acid limitation leads to the stabilization of many transcripts independent of (pp)pGpp (Geiger *et al*., 2010). Thus, from these analyses, it is difficult to draw firm conclusions on the primary transcriptional changes imposed by (pp)pGpp synthesis. Recently, one study tried to circumvent this drawback by transcriptional induction of (pp)pGpp synthetase in *E. coli* and gained major new insights (Sanchez-Vazquez *et al*., 2019).

Here, we aimed to compare the RSH-, RelQ- and RelP-mediated effects on nucleotide pools, transcription and functional consequences without imposing nutrient starvation. Therefore, the synthetase domain of RSH (RSH-Syn), RelP and RelQ were expressed from an anhydrotetracycline (ATc)-inducible promoter in a (pp)pGpp^0^ strain in which the enzymatic domains of all three synthetases were deleted. Through RNA-Seq analyses, we identified new (pp)pGpp-regulated genes, many of which are involved in the oxidative stress response, iron storage and the synthesis of phenol-soluble modulins (PSMs). Thus, (pp)pGpp synthesis contributes not only to PSM-derived ROS production but also to protection from these toxic molecules.

## Results

### Changes in the nucleotide pools after transcriptional induction of RSH-Syn and RelQ

We first compared the stringent response imposed by mupirocin (tRNA synthase inhibitor) with the genetic induction of (pp)pGpp synthetases. RelQ or RSH-Syn (N-terminal part of RSH in which the hydrolase domain was mutated) were expressed using an ATc-inducible expression system in a pppGpp^0^ strain background. Strain (pp)pGpp^0^ is unable to synthesize (pp)pGpp due to mutations in all three (pp)pGpp synthetases (full deletion of *rsh*, synthetase mutation in *relP* and *relQ*). Strains were grown to an early exponential growth phase and gene expression was induced for 30 min. Consistent with previous results (Kastle *et al*., 2015), treatment of the (pp)pGpp^0^ strain with mupirocin resulted in a significant increase in the GTP pool (Fig.1). Induction of the stringent response in the wild type by either mupirocin or transcriptional induction of RSH-Syn resulted in similar changes in the nucleotide pools: an immediate increase in the ATP pool, a decrease in the GTP pool and synthesis of the alarmones pppGpp, ppGpp, and pGpp (Fig. 1). After induction of RelQ, only minor changes in the nucleotide pool were detectable. Thus, the effect of RelQ on the nucleotide pools was significantly lower than that of RSH-Syn.

**Fig. 1.**
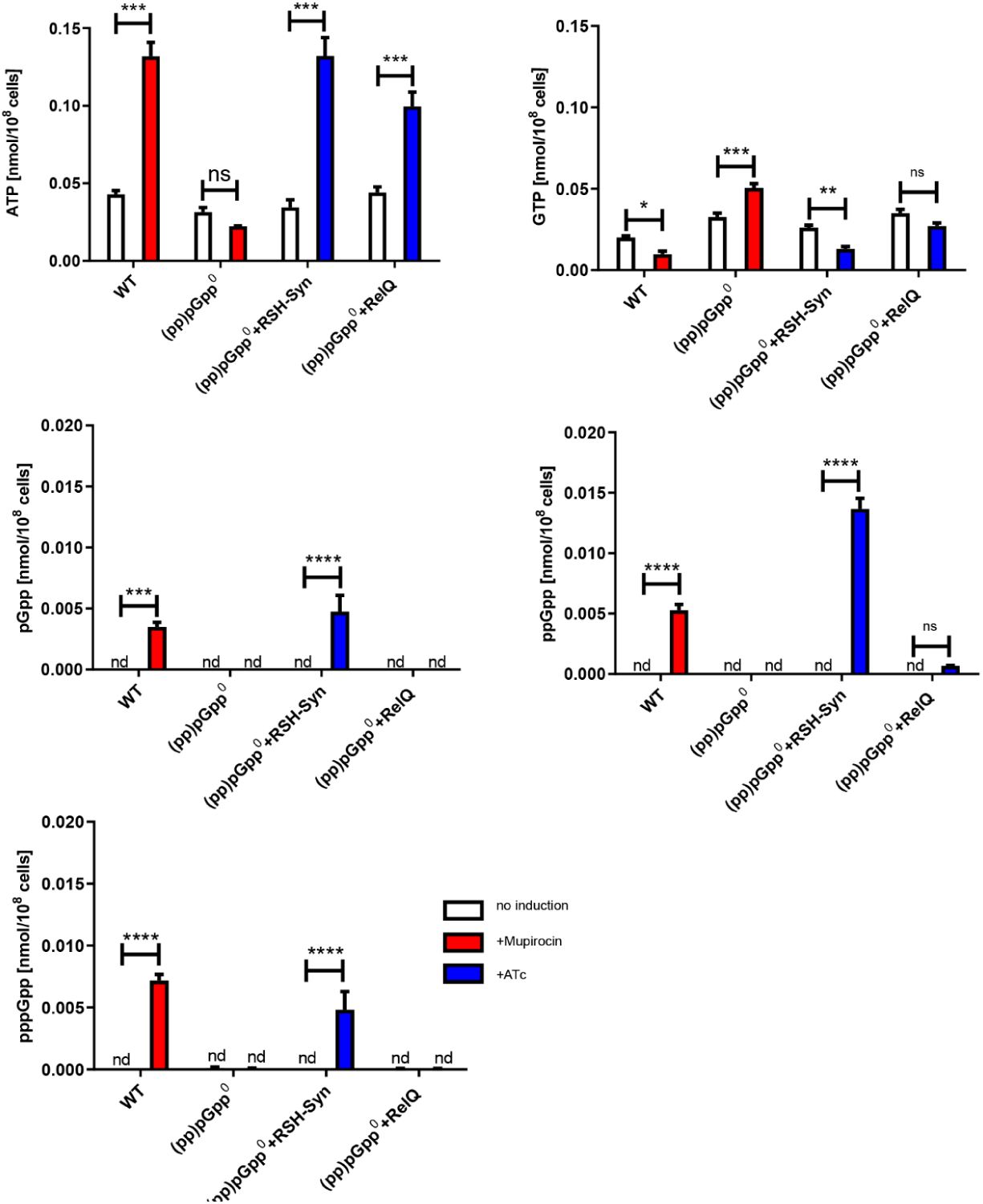
Changes in the nucleotide pool after mupirocin treatment or transcriptional induction of RSH-Syn or RelQ. Strain HG001 and derivatives were grown to OD_600_ = 0.3 and treated for 30 min with or without 0.125 µg/ml mupirocin (red) or 0.1 µg/ml ATc (blue). Nucleotide analyses were performed using mass spectrometry (ESI-TOF) in negative ion mode. Error bars represent SEM (n=3) from three biological replicates. Statistical significance was determined by two-way ANOVA with Tukey’s posttest, *p ≤ 0.05, **p ≤ 0.01, ***p ≤ 0.001 and ****p ≤ 0.0001.

### Impact of (pp)pGpp synthesis on the transcriptome

We next analyzed the impact of (pp)pGpp synthesis after induction of RSH-Syn or RelQ on mRNA abundance. RNA-Seq data revealed that 1449 genes and sRNAs were significantly affected by either RSH-Syn (total: 1388, 717 up, 671 down) or RelQ (total: 352, 223 up, 129 down). The expression of most of the RelQ-affected genes was also changed in response to RSH-Syn induction (Fig. 2A. Suppl. Tab. 1). However, consistent with the nucleotide measurements (Fig. 1), the effect of RelQ induction was less prominent (Suppl. Tab 1). We compared the data with previous microarray analyses after induction of the stringent response imposed by amino acid limitation (Geiger *et al*., 2012). Most of the previously identified stringent response genes were confirmed by the RNA-Seq analysis (Fig. 2A). Of note, in the present analysis, only genes with at least three-fold differences and a significance level of p < 0.001 were included in the analysis shown in Fig. 2 and Suppl. Tab. 1. Therefore, some of the previously detected genes by Geiger et al., (Geiger *et al*., 2012) were excluded, although most of them showed the same tendency (see Suppl. Tab. 2). Despite the higher stringency in the analysis, the present analysis revealed far more (pp)pGpp-regulated genes and sRNAs. Genes were classified into functional categories using the SEED annotation (http://pubseed.theseed.org). (pp)pGpp induction resulted in the downregulation of many genes involved in protein, RNA and DNA metabolism, consistent with previous results that the stringent response mainly leads to the shutdown of translation and replication (Geiger *et al*., 2012) (Fig. 2B). More than 500 RNAs were significantly upregulated upon RSH-Syn induction (Suppl Tab. 1). Most of them were sRNAs or coded for hypothetical proteins with unknown function. Many of the amino acid biosynthesis gene clusters were also upregulated. Most of them were part of the CodY regulon and thus likely regulated by lowering of the GTP pool. Phage-encoded genes were also upregulated, indicating phage-inducing conditions. This is in line with the upregulation of *recA* and *lexA*. We next sorted for genes that were most affected by RSH-Syn induction (Fig. 2C, Suppl. Tab. 1). Many of the genes were assigned to iron acquisition/metabolism (upregulation of genes involved in iron storage; downregulation of genes involved in siderophore biosynthesis and iron transport), stress response (*dps, sodA, katA ahpC, uspA1/2, asp23, ptpA*, and *msrA2)*, and virulence (upregulation of *psmsα/ß*; downregulation of *agr*). For further analysis, we used *ftnA, dps, agr* and *psm*α as read-outs for (pp)pGpp-mediated activities under various conditions.

**Fig. 2.**
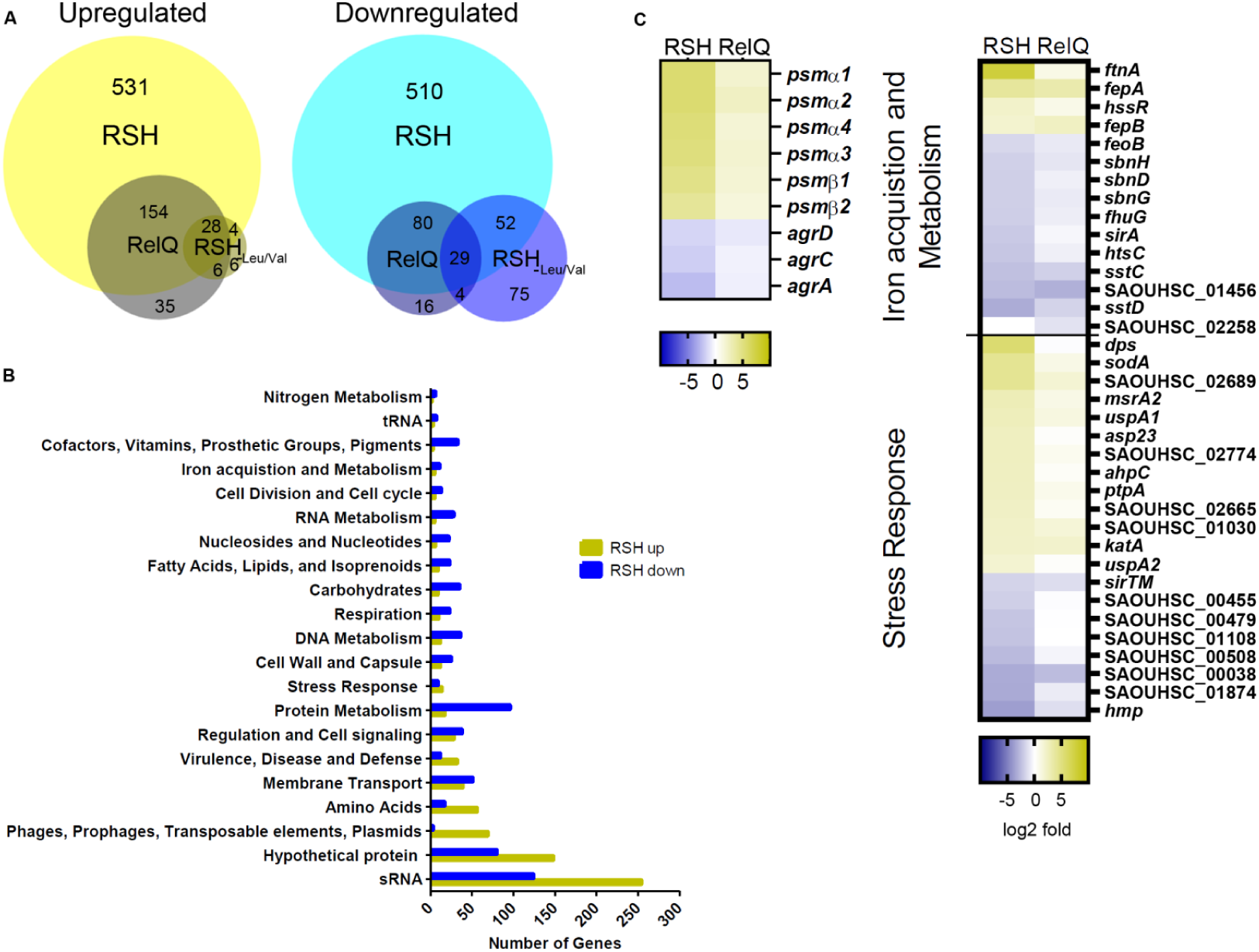
Global changes in gene expression upon transcriptional induction of RSH-Syn or RelQ. pppGpp^0^ with inducible RSH-Syn or relQ was grown to OD600 = 0.3 and treated for 30 min without or with 0.1 µg/ml ATc. **A**. Number of genes or sRNAs upregulated (yellow) or downregulated (blue) after induction in comparison to uninduced cultures (< 3-fold difference, p< 0.001). Previously, described stringent genes (Geiger *et al*., 2012) are indicated as RSH_-Leu/Val_. **B**. Genes with significant changes after induction of RSH-Syn (< 3-fold difference, p< 0.001) according to functional categories. **C**. Heatmap representing RSH-Syn-dependent up- and downregulated genes assigned to the functional categories iron acquisition and metabolism, stress response and Agr-related genes.

### Comparison of the mupirocin-induced stringent response and transcriptional induction of RSH-Syn

We compared the expression of the selected genes after induction of the stringent response via mupirocin and after transcriptional induction of RSH-Syn. We verified the upregulation of *ftnA, dps* and *psm* under both conditions (Fig. 3A). Mupirocin also resulted in *ftnA* and *dps* activation in the (pp)pGpp^0^ strain, although to a lesser extent, indicating additional (pp)pGpp-independent effects of mupirocin on the expression of these genes. The activation of *psm* expression by (pp)pGpp is clearly not correlated to *agr* expression. Agr is the main activator required for *psms* expression (Queck *et al*., 2008). Notably, the expression of the *agr* operon was even lower upon (pp)pGpp synthesis (Suppl. Tab. S1 and Fig. 3A), indicating that (pp)pGpp induces *psm* expression independent of Agr.

**Fig. 3.**
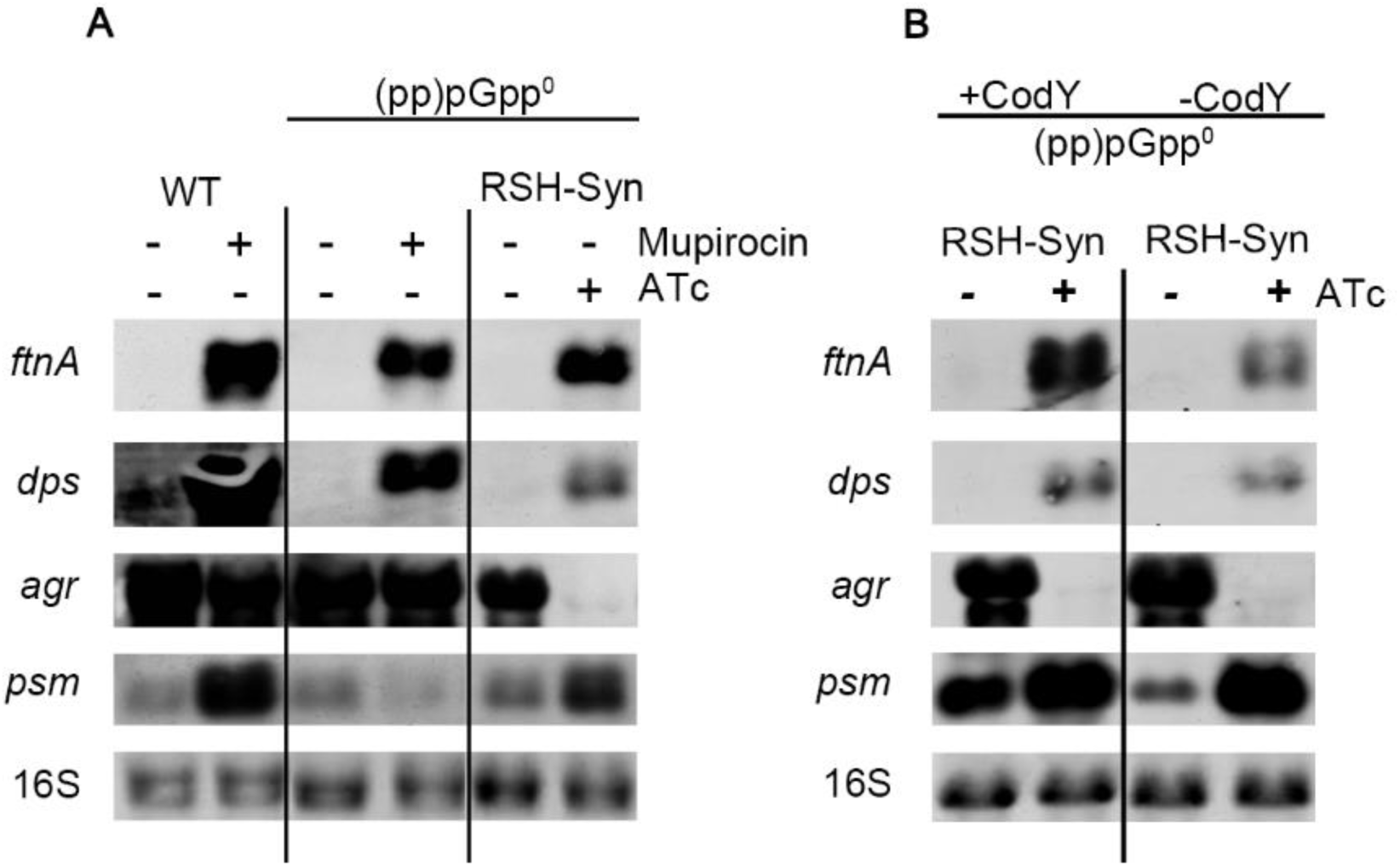
Correlation of mupirocin-induced stringent response and transcriptional induced RSH-Syn of selected CodY-independent genes. Strain HG001 and derivatives were grown to OD_600_ = 0.3 and treated for 30 min with or without 0.125 µg/ml mupirocin or 0.1 µg/ml ATc (mutant strains with inducible RSH-Syn). For Northern blot analysis, RNA was hybridized with digoxigenin-labelled probes specific for *ftnA, dps, psm* or *agrA*. The 16S rRNA detected in ethidium bromide-stained gels is indicated as a loading control in the bottom lane.

### CodY-independent activation of gene expression by RSH-Syn induction

(pp)pGpp synthesis leads to the lowering of the GTP pool and subsequently to de-repression of CodY target genes. Indeed, many of the genes that were upregulated in response to RSH-Syn induction belong to the CodY regulon (Suppl. Tab. 1). However, none of our selected marker genes are known to be regulated via CodY. To exclude CodY-dependent regulation, we compared the expression of the selected marker genes in a *codY*-negative mutant. Induction of RSH-Syn or RelQ resulted in similar expression patterns in *codY*-positive and *codY*-negative backgrounds (Fig. 3B and Fig. S2 for relQ). This shows that (pp)pGpp impacts the expression of these genes independent of CodY.

### RSH-Syn induction influences the oxidative stress response and virulence independent of PerR, Fur or SarA

Some of the prominent (pp)pGpp-activated genes are known to be under the control of other global regulators, such as PerR, Fur and SarA (Gaupp *et al*., 2012). *ftnA, dps, ahpC* and *katA (*Suppl. Tab. 1) are likely controlled via PerR binding to a conserved PerR-binding motif based on the public databases RegPrecise (Novichkov *et al*., 2013) and Aureowiki (Fuchs *et al*., 2018). We speculated that (pp)pGpp induction of these genes may somehow be mediated via PerR activity. Therefore, we induced RSH-Syn in a *perR/*(pp)pGpp^0^ background (Fig. 4A). As expected, *ftnA* and *dps* were both upregulated in the *perR* mutants. Inducing RSH-Syn showed an even higher upregulation of *ftnA*, indicating that (pp)pGpp acts in addition and independent of PerR. For *dps*, the *perR* mutation alone resulted in high expression, which was not further increased by (pp)pGpp, indicating that *dps* is expressed at its maximum level in the *perR* mutant. *PerR* deletion resulted in a slight decrease in *psm* expression, which was compensated by RSH-Syn induction. Thus, (pp)pGpp also affects gene expression in a *per*-negative background.

**Fig. 4.**
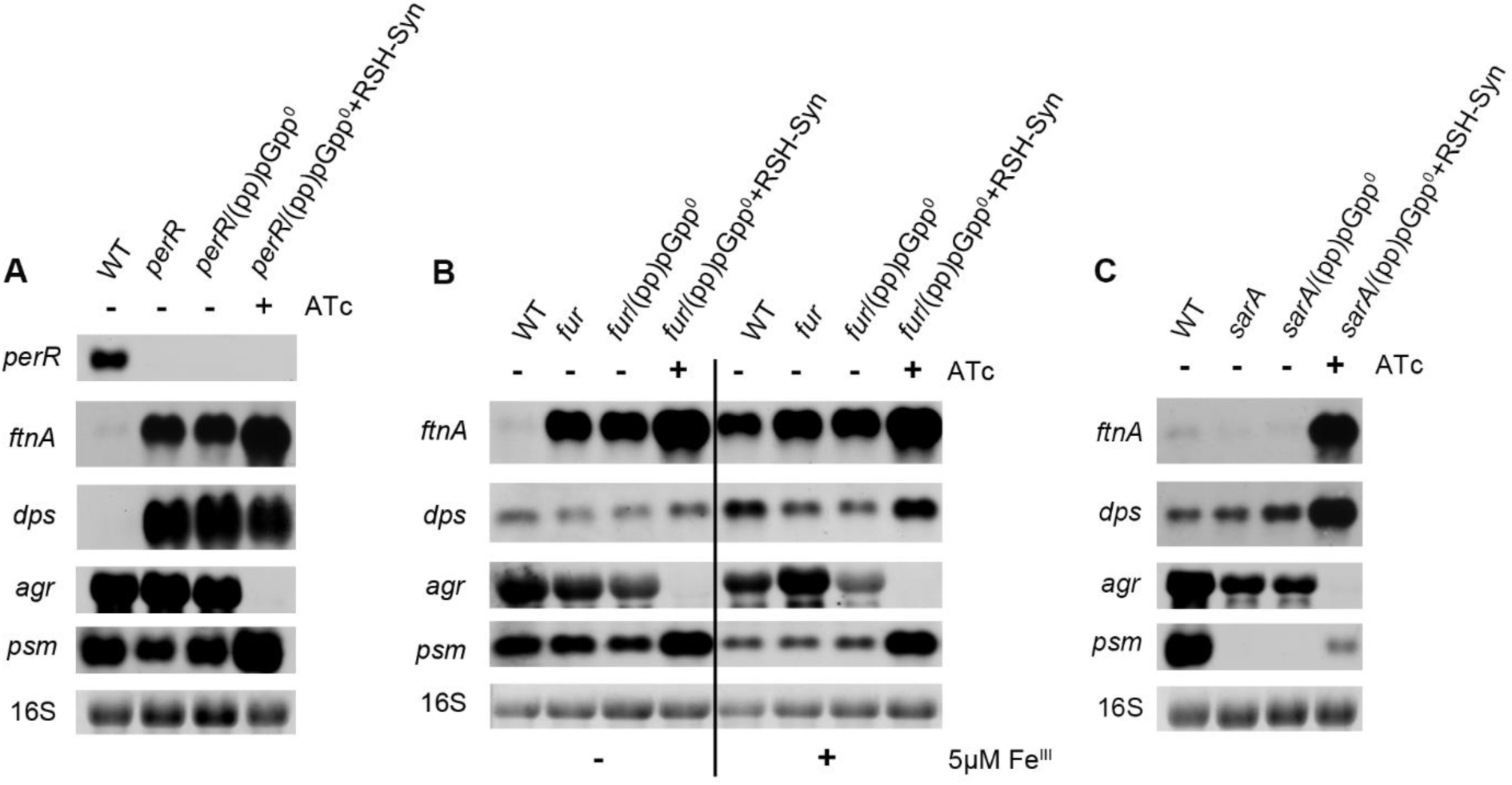
(pp)pGpp-dependent transcriptional changes are independent of PerR, Fur or SarA. Strain HG001 and derivatives were grown to OD_600_ = 0.3 and treated for 30 min without or with 0.1 µg/ml ATc (mutant strains with inducible RSH-Syn). For Northern blot analysis, RNA was hybridized with digoxigenin-labelled probes specific *for ftnA, dps, psmα* or *agrA*. The 16S rRNA detected in ethidium bromide-stained gels is indicated as a loading control in the bottom lane.

We found that many of the genes affected by RSH-Syn are indicative of iron overload conditions (e.g. upregulation of *ftnA* and *dps*). We asked whether the iron-responsive regulator Fur is involved in the regulation of these genes. Therefore, we induced RSH-Syn in a *fur*/(pp)pGpp^0^ background under low and high iron conditions (Fig. 4B). Independent of the availability of iron, *ftnA, dps* and *psm* were upregulated and *agr* was downregulated after RSH-Syn induction in the *fur*-negative background.

SarA was shown to activate transcription of the *agr* operon (Heinrichs *et al*., 1996, Zielinska *et al*., 2011) and proposed to be involved in oxidative stress sensing via a single Cys9 residue (Sun *et al*., 2012, Ballal & Manna, 2010, Grosser *et al*., 2016). *sarA* was found to be significantly upregulated by RSH-Syn (Suppl. Tab. 1). To analyze whether SarA is involved in (pp)pGpp regulation, we induced RSH-Syn in a *sarA*/(pp)pGpp^0^ background (Fig. 4C). *ftnA* and *dps* expression was not influenced by *sarA* mutation. *agr* and *psm* expression was downregulated in the *sarA* mutant, consistent with the proposed activation of the *agr* system by SarA (Heinrichs *et al*., 1996). Inducing RSH-Syn again showed the typical induction of *ftnA, dps* and *psm* and repression of *agr* in the *sarA* mutant. Taken together, our results show that (pp)pGpp regulates the selected genes independent of the main transcriptional regulators (CodY, PerR, Fur and SarA) known to control the expression of some of them.

### Effects of RSH-Syn induction in strain USA300

Thus far, we concentrated our analysis on the effects of RSH-Syn induction in strain HG001. We also constructed a (pp)pGpp^0^ mutant in strain USA300 and analyzed gene expression after RSH-Syn induction (Fig. 5A). As in HG001, induction of RSH-Syn resulted in the typical induction of *ftnA, dps* and *psm* and downregulation of *agr* in strain USA300.

**Fig. 5.**
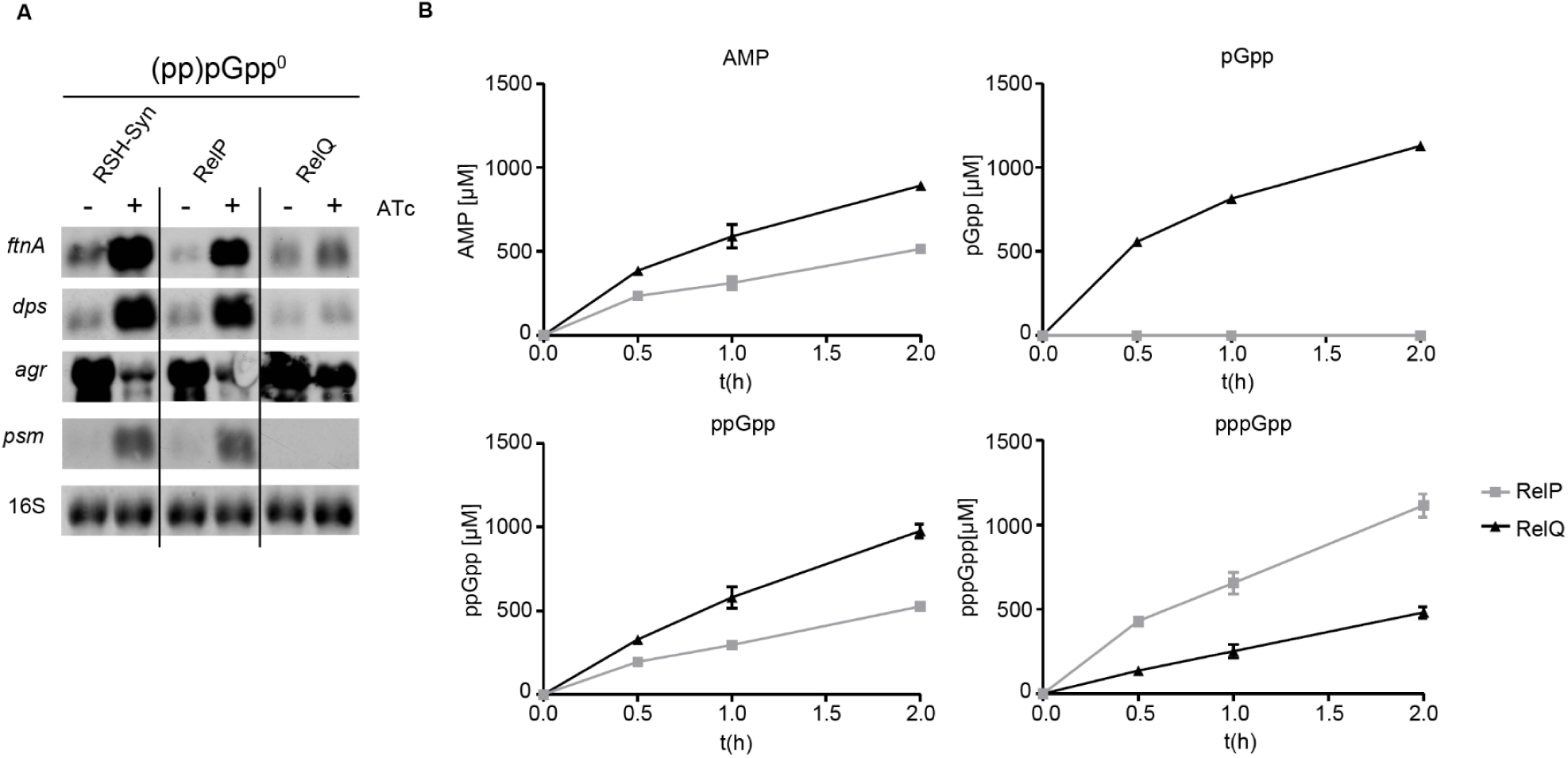
RelP activity *in vitro* and *in vivo*. **A**. Strain USA300 and derivatives were grown to OD_600_ = 0.3 and treated for 30 min without or with 0.1 µg/ml ATc (mutant strains with inducible RSH-Syn, RelP or RelQ). For Northern blot analysis, RNA was hybridized with digoxigenin-labelled probes specific for ftnA, dps, psmα or agrA. The 16S rRNA detected in ethidium bromide-stained gels is indicated as a loading control in the bottom lane. **B**. Purified RelP and RelQ (0.2 µM) were incubated with 1 mM ATP, GMP, GDP and GTP at 37°C for 30 min, 1 h and 2 h. The products AMP and (pp)pGpp were quantified by mass spectrometry.

### RelP induction is similar to RSH-Syn induction

The RNA-Seq analysis revealed that induction of RelQ had only a minor effect on the target genes compared to induction of RSH-Syn, although RelQ was highly expressed after ATc treatment (Suppl. Tab. 1). We analyzed whether this was also true in strain USA300 and whether induction of the homolog enzyme RelP would be similar to induction of RelQ. We verified that induction of RelQ had only minor effects on marker gene expression (Fig. 5A). However, induction of RelP was highly effective, resulting in an expression pattern comparable to induction of RSH-Syn.

Since we observed a severe difference between induction of RelP and RelQ *in vivo*, we wondered whether RelP is simply a more active enzyme. Therefore, we measured (pp)pGpp synthesis of recombinant RelP and RelQ *in vitro* (Fig. 5B). RelP or RelQ (0.2 µM) were incubated with ATP and an equal molar mixture of the potential substrates GTP, GDP and GMP. RelQ was even more active than RelP, as indicated by increased levels of generated AMP. However, RelP preferentially synthesizes pppGpp, whereas RelQ preferentially synthesizes ppGpp and pGpp.

### (pp)pGpp is involved in oxidative stress resistance

PSMs were shown to result in intracellular production of reactive oxygen species (ROS) (George *et al*., 2019). Thus, it is likely that under stringent conditions, (pp)pGpp-mediated PSM synthesis further increases ROS formation. One might speculate that PSM-mediated ROS production triggers the expression of the oxidative stress genes detected in the transcriptome analysis. In this case, RSH-Syn induction should not result in the induction of these genes under anaerobic conditions, where ROS cannot be produced. However, RSH-Syn induction resulted in the same transcriptional pattern regardless of whether bacteria were grown with or without oxygen (Fig. 6A). Thus, (pp)pGpp-mediated gene alterations of the selected marker genes are not a consequence of ROS generation by PSMs.

**Fig. 6.**
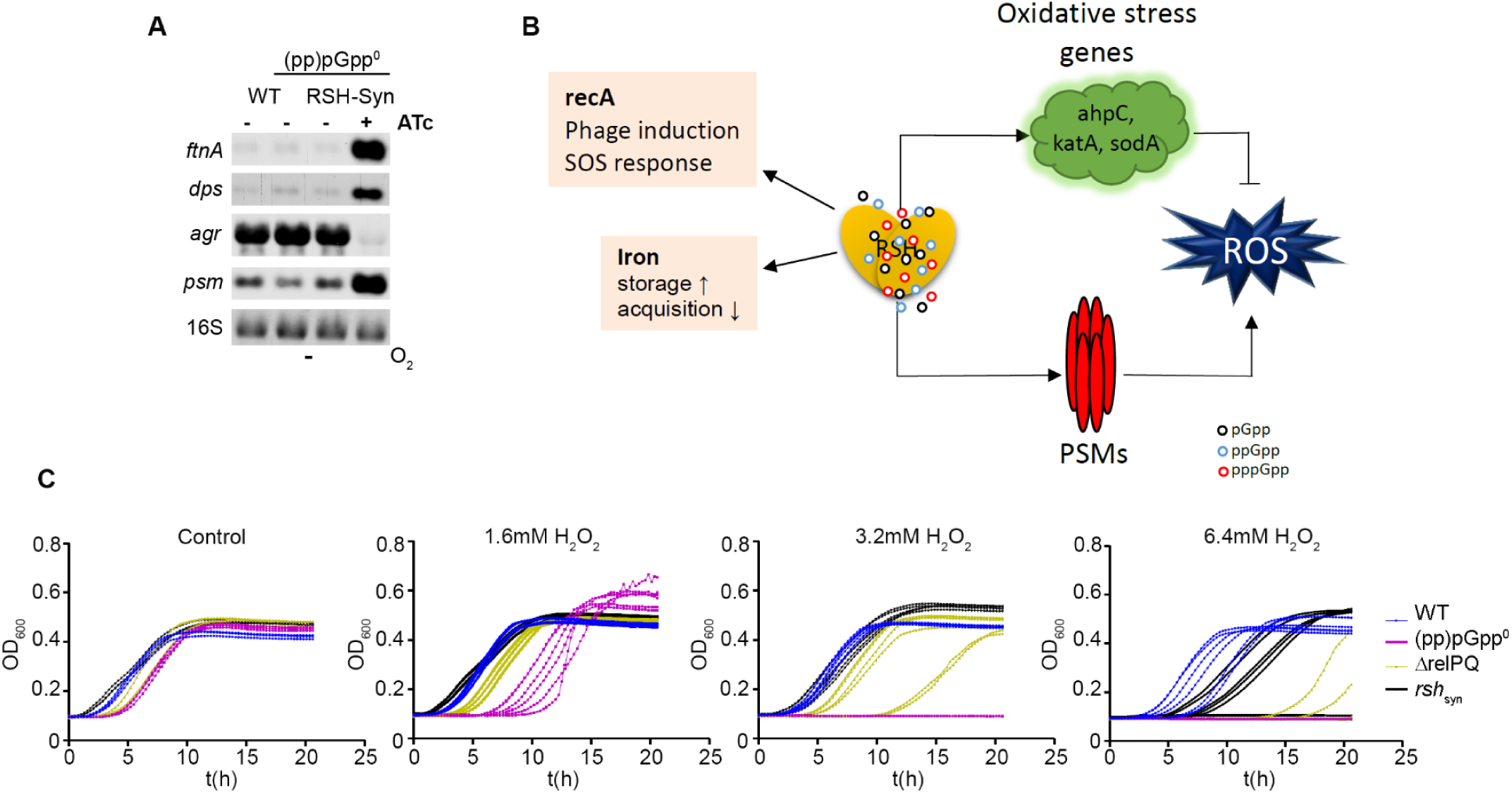
Functional link between stringent response and oxidative stress. **A**. Strain HG001 and derivatives were grown anaerobically to OD_600_ = 0.3 and treated for 30 min without or with 0.1 µg/ml ATc (mutant with inducible RSH-Syn). For Northern blot analysis, RNA was hybridized with digoxigenin-labelled probes specific for *ftnA, dps, psm* or *agrA*. The 16S rRNA detected in ethidium bromide-stained gels is indicated as a loading control in the bottom lane. **B**. (pp)pGpp leads to upregulation of oxidative stress genes. These upregulated genes are beneficial to counteract endogenous (PSMs) or exogenous (e.g., H_2_O_2_) ROS. Upregulation of SOS and phage genes might be a consequence of ROS accumulation, e.g., by PSMs. Upregulation of iron storage proteins may be beneficial to protect bacteria from ROS generated by the Fenton reaction. **C** WT, (pp)pGpp^0^, Δ*relPQ* and *rel*_syn_ mutants were diluted from overnight culture to an OD = 0.1 and challenged with different H_2_O_2_ concentrations, and growth was monitored over time

These data indicate that (pp)pGpp simultaneously activates ROS-producing PSMs as well as ROS defence systems to prepare cells to withstand oxidative stress (Fig. 6B). To verify this hypothesis, we challenged wild-type and mutant strains deficient in (pp)pGpp synthesis with H_2_O_2_. The (pp)pGpp^0^ strain was indeed more sensitive to oxidative stress (wild type MIC = 6.4 mM H_2_O_2;_ (pp)pGpp^0^ MIC = 3.2 mM H_2_O_2_). Under these non-induced conditions, (pp)pGpp might be derived from any of the pppGpp synthetases. We also analyzed a *relPQ* mutant and a *rsh-syn* mutant in which the synthetase domain of RSH was mutated. Both strains showed an intermittent phenotype in which the MIC varied between 3.2 and 6.4 mM when biological replicates were analyzed. To follow up on these ambiguities, we monitored growth after the addition of H_2_O_2_ (Fig. 6C). There was high variation in the lag time between biological replicates. Replicates of the *relPQ* or *rsh-syn* mutant showed a delayed lag phase, and some of the replicates could not grow. The delay of the lag phase was more prominent for the *relPQ* mutant than for the *rsh-syn* mutant. Nevertheless, none of the (pp)pGpp^0^ replicates could resume growth, and this result was consistent with the reproducible lowered MIC of this strain, indicating that (pp)pGpp indeed protects against oxidative stress.

## Discussion

We chose a genetic approach to define the early transcriptional response upon (pp)pGpp synthesis without the need to apply additional stress conditions. As expected from previous studies, (Geiger *et al*., 2012, Reiss *et al*., 2012) (pp)pGpp synthesis resulted in severe downregulation of the translational machinery and de-repression of CodY target genes. Many additional (pp)pGpp-regulated genes and sRNAs that are presumably important for the survival of *S. aureus* during starvation conditions were identified. Here, we focused mainly on genes that were found to be activated upon (pp)pGpp synthesis in a CodY-independent manner, particularly *psm, ftnA* and *dps*. The (pp)pGpp-dependent activation of these genes can also occur in strains missing the prototypic proteinaceous transcriptional regulators PerR, Fur, or SarA. These regulators are well known to be involved in the regulation of the selected genes. However, they need to be activated through oxidative stress and/or iron (Gaupp *et al*., 2012). Thus, (pp)pGpp functions as a complementary, immediate message, allowing cells to react to adverse conditions such as amino acid starvation or cell wall stress. Under these conditions, upcoming oxidative stress seems to be anticipated, and (pp)pGpp prepares the cells for survival, e.g. ROS challenges. Indeed, a (pp)pGpp^0^ strain is more sensitive to H_2_O_2._ Both RSH and RelP/RelQ contribute to the protective effect.

### (pp)pGpp leads to *psm* activation

One of the most prominent effects of (pp)pGpp synthesis is the upregulation of *psmα and psmβ*, confirming previous microarray analyses (Geiger *et al*., 2012). PSMs are a family of amphipathic, alpha-helical peptides that have multiple roles in staphylococcal pathogenesis and contribute a large extent to the pathogenic success of virulent staphylococci (Cheung *et al*., 2014, Peschel & Otto, 2013). (pp)pGpp-dependent *psm* expression within neutrophils was shown to be crucial for survival after phagocytosis (Geiger *et al*., 2012). However, PSMs also interact with the membrane of the producer, promote the release of membrane vesicles from the cytoplasmic membrane via an increase in membrane fluidity (Schlatterer *et al*., 2018, Wang *et al*., 2018), reduce persister formation (Bojer *et al*., 2018, Xu *et al*., 2017) and are involved in self-toxicity via ROS formation (George *et al*., 2019). Interestingly, *psm* activation was not correlated with the activation of the quorum sensing Agr system, the main regulator required for *psm* expression (Queck *et al*., 2008). Agr was even repressed under (pp)pGpp-inducing conditions. Previously, analysis of a clinical isolate overproducing (pp)pGpp also indicated that (pp)pGpp leads to *agr* inhibition (Gao *et al*., 2010). Thus, (pp)pGpp-mediated *psm* activation is clearly uncoupled from *agr* expression. Recently, the sRNA Teg41 (S131) (Zapf *et al*., 2019) and the transcriptional regulator MgrA (Jiang *et al*., 2018) were found to interfere with *psm* expression. However, it is unlikely that they mediate the (pp)pGpp regulatory effect because transcription of these regulators was unaltered based on our RNA-Seq analysis (Suppl. Tab. S1). Thus, the molecular mechanism by which (pp)pGpp leads to *psm* activation has to be elucidated. It is likely that the accompanying changes in the ATP/GTP ratios are crucial for this activation pattern. *psm* promoters might be sensitive to the concentration of the initiating nucleoside triphosphate (iNTP). The +1 position (e.g., G or A) dictates whether transcriptional initiation/elongation requires high GTP or ATP levels, respectively (Krasny *et al*., 2008). Various sequence combinations determine whether a promoter is sensitive to iNTP (Sojka *et al*., 2011). Such sequence motifs are hard to predict within *psm* promoters. However, both the p*smα* and *psmß* operons start with an A at the +1 position (Queck *et al*., 2008), which might explain the higher expression due to the increased ATP levels following (p)ppGpp synthesis.

### (pp)pGpp and oxidative stress response

Genes whose expression is indicative of iron overload conditions were also highly affected by (pp)pGpp. Recently, a similar effect was reported for *Vibrio cholera* (Kim *et al*., 2018). Here, the expression of the iron transporter FbpA was repressed via (pp)pGpp, resulting in a reduction of intracellular free iron required for the ROS-generating Fenton reaction. This contributed to reducing antibiotic-induced oxidative stress and thus tolerance, and it is likely that this is also the case in *S. aureus*. In addition to interfering with iron metabolism, other genes involved in oxidative stress were activated by (pp)pGpp. A link between the stringent response and oxidative stress response has been observed in different organisms, although the underlying mechanisms and outcome might be highly diverse. (pp)pGpp-dependent upregulation of superoxide dismutase (SOD) was described in *B. suis* (Hanna *et al*., 2013) and *P. aeruginosa* (Martins *et al*., 2018). SOD was shown to be the key factor responsible for (pp)pGpp-mediated multidrug tolerance in *P. aeruginosa* (Martins *et al*., 2018). Moreover, (pp)pGpp-deficient strains are often found to be more sensitive to oxidative stress (Yan *et al*., 2009, Holley *et al*., 2014, Wang *et al*., 2016). However, in *E. faecalis*, a (pp)pGpp^0^ strain grew faster and to a higher growth yield than its parent in the presence of H_2_O_2_ (Abranches *et al*., 2009).

Here, we show that the stringent response in *S. aureus* leads to the activation of ROS-inducing toxins and simultaneous expression of the detoxifying system to protect the producer. This is likely a special advantage for the pathogen once it encounters neutrophils and elevated ROS. (pp)pGpp dependent PSM synthesis is required to escape from within cells after phagocytosis (Surewaard *et al*., 2013, Geiger *et al*., 2012). The upregulation of the oxygen stress programme will help protect the cell from endogenous as well as exogenous ROS.

### Comparison of RSH-Syn, RelQ and RelP activity

We compared the activity of RSH-Syn, RelQ and RelP. Nucleotide profiling as well as transcriptional analysis showed that induction of RelQ results in nucleotide changes similar to induction of RSH-Syn, although to a much lesser extent. In contrast, RelP was equally active as RSH-Syn. Thus, RelQ activity seems to be restricted *in vivo* under our growth conditions. Comparison of RelP and RelQ from other organisms revealed that RelQ is inhibited through RNA binding and auto-activated by (pp)pGpp (Manav *et al*., 2018, Steinchen *et al*., 2018). We analyzed RelQ activity in an (pp)pGpp background under non-stress conditions where RelQ activity is likely restricted via RNA binding and/or the missing basal (pp)pGpp provided by other synthetases. The conditions that would relieve this restriction remain to be determined. Analysis of purified RelP and RelQ revealed that both enzymes are equally active. The most striking difference was that RelQ can use GMP as a substrate to produce pGpp, whereas no pGpp synthesizing activity was detectable for purified RelP. pGpp synthesis was already described for *Corynebacterium glutamicum* (Ruwe *et al*., 2017) or RelQ from *E. faecalis* (Gaca *et al*., 2015b) Of note, we also detected RSH-dependent pGpp synthesis via either mupirocin treatment or RSH-Syn induction *in vivo*. pGpp was shown to exert inhibitory effects similar to ppGpp for enzymes involved in GTP biosynthesis (Gaca *et al*., 2015b). Previous nucleotide analyses in *S. aureus* (Crosse *et al*., 2000) and other organisms (Pao & Gallant, 1979) revealed another molecule and proposed to be even more active than pppGpp regarding inhibition of some target proteins (Pao & Dyess, 1981). GTP, pGpp and ppGp are difficult to differentiate due to their identical molecular weights, and the biological relevance of these molecules remains to be elucidated. Nevertheless, the function of the two SAS enzymes often present together within one organism indicates that they might fulfil distinct functions. RelQ requires posttranscriptional activation and preferentially synthesizes pGpp. The different ratios of the alarmones pppGpp, ppGpp and pGpp may contribute to small differences in the stringent response outcome (Mechold *et al*., 2013). However, the overall changes in the nucleotide pool and transcriptional changes between the three (pp)pGpp synthetases of *S. aureus* are largely similar.

## Materials and Methods

### Strains and growth conditions

Strains and plasmids are listed in Suppl. Tab. S3. For strains carrying a resistance gene a concentration of 10 µg/ml chloramphenicol, 10 µg/ml erythromycin or 100 µg/ml ampicillin was used only for overnight cultures. *S. aureus* strains were grown overnight in chemical defined medium (CDM) (Pohl *et al*., 2009), diluted to an optical density (OD_600_) of 0.05 and grown until the early exponential phase OD_600_ = 0.3 with shaking (220 rpm, 37°C). Gene expression in strains carrying a plasmid with an ATc-inducible promoter was induced at OD_600_ = 0.3 with 0.1 µg/ml ATc for 30 min. For anaerobic growth the strains were diluted to an OD_600_ = 0.05 in hungate tubes (Chemglass), completely filled with CDM. ATc was applied using a syringe at OD_600_ = 0.03. For OD measurements and RNA isolation, aliquots were drawn with a syringe.

### Generation of (pp)pGpp^0^ mutant in USA300 JE2

For the USA300 (pp)pGpp^0^ mutant (USA300-229-230-263), lysates were prepared from RN4220 strains containing the mutagenesis vectors pCG229, pCG230 and pCG263, respectively (Suppl. Tab S3). After plasmid transduction of USA300 JE2, mutagenesis was performed as previously described (Bae & Schneewind, 2006). To avoid toxic accumulation of (pp)pGpp the genes were mutated in the order *relP, relQ* and finally *rsh*. Mutations were verified by PCR using oligonucleotides listed in Suppl. Tab. S4.

### Generation of *perR, fur, sarA* and *psm*α/β (pp)pGpp^0^ mutants in HG001

Φ11 lysates were generated from transposon mutants NE665 (*perR*), NE99 (*fur*) and NE1193 (*sarA*) from the NARSA transposon library (Fey *et al*., 2013) to transduce *S. aureus* strains HG001 and (pp)pGpp^0^. *psm*α or *psmβ* mutations were transduced using Φ11 lysates from previously described mutants (Geiger *et al*., 2012). All transductants were verified by PCR using oligonucleotides listed in Suppl. Tab. S4.

### RNA isolation and Northern Blot analysis

RNA isolation and northern blot analysis were performed as described previously (Goerke *et al*., 2000). Briefly, bacteria were pelleted and resuspended in 1 ml TRIzol (Thermo Fisher Scientific) and lysed using zirconia/silica beads (0,1mm diameter) and a high speed homogenizer. RNA was isolated following the recommended procedure by TRIzol manufacturer. For RNA-Seq analysis RNA from the aqueous phase was further purified following the RNA-isolation protocol by Amp Tech ExpressArt^®^ RNA ready. Transcripts on the Northern blot were detected by dioxigenin-labeled probes, which were generated by a DNA-labelling PCR-Kit (Roche Life Science).

### Purification of RSH-Syn, RelP and RelQ

Proteins were purified as previously described (Gratani *et al*., 2018, Geiger *et al*., 2014). Briefly, plasmids carrying RelP or RelQ were freshly transformed into *E. coli* B21 and grown for 16 hours at RT in LB supplemented with D(+)-lactose-monhydrate (12.5 g/l) and 100 µg/ml ampicillin. Bacteria were harvested (20 min, 3000 x g, 4 °C) and resuspended in ice-cold low-KCl buffer A (20 mM HEPES (pH 7.4), 200 mM NaCl, 20 mM MgCl2, 20 mM KCl, 30% (v/v) glycerol, and 40 mM imidazole) supplemented with 10 μg/ml DNAse and cOmpleteTM protease inhibitor cocktail (Roche). Bacteria were lysed by a french press at 1000 psi and centrifuged (50,000 x g, 45 min, 4 °C). The clear supernatant was filtered (0.22-μm pore size) and loaded onto a 1-ml HisTrap HP column (GE Healthcare Life Sciences) equilibrated with low-KCl buffer A. Proteins were purified by an ÄKTA purification system (GE Healthcare Life Sciences) and eluted with an imidazole gradient to a final concentration of 500 mM. The eluted fractions were loaded on SDS-PAGE, protein-containing fractions were concentrated by an Amicon Ultracel-30K ultracentrifugal device. Size exclusion columns was performed (HiLoad 16/600 Superdex 200 pg, GE Healthcare Life Sciences) using ice-cold low-KCl SEC buffer (20 mM HEPES (pH 7.0), 200 mM NaCl, 20 mM MgCl2, 20 mM KCl, and 30% (v/v) glycerol). Protein containing fractions were concentrated by ultrafiltration and stored at −80°C.

### *In vivo* nucleotide extraction

Nucleotids were isolated based on published protocol (Juengert *et al*., 2017). Briefly, strains were grown in CDM overnight, diluted to an OD_600_ = 0.05 and grown in CDM until an OD_600_ = 0.3. Strains were split and treated with or without 0.1 µg/ml ATc for 30 min at 37°C and 220 rpm shaking. 100 ml bacterial cultures were harvested and transferred into 50 ml centrifuge tubes half filled with ice and centrifuged (5 min, 5000 x g, 4°C). Pellets were immediately frozen in liquid nitrogen and stored at −80°C until usage. Samples were thawed on ice and resuspended in 2M formic acid and incubated for 30 min. Resuspended bacteria were lysed by high speed homogenizer using zirconia/silicia beads (0.1 mm diameter) and kept on ice for 30 min. The aqueous phase was collected and mixed with 3 ml 50 mM NH_4_OAc (pH 4.5), loaded on columns (OASIS Wax cartridge 3xcc) and centrifuged (5000 x g, 5min, 4°C). Columns were pre-treated first with pure 3 ml methanol and then with 3 ml 50 mM NH_4_OAc (pH 4.5). Samples were washed first with 3 ml 50 mM NH_4_OAc (pH 4.5) followed by a washing step with 3 ml methanol. Elution was performed with 1 ml of 20% methanol, 10% NH_4_OH. Eluted nucleotides were flash frozen in liquid nitrogen and lyophilized overnight. Lyophilized nucleotides were resuspended in 100 µl ddH_2_O and analyzed via HPLC-MS. pppGpp and ppGpp standard molecule were purchased by. pGpp was synthesized starting from conveniently protected guanosine and employing both phosphoramidite and phosphotriester methods (Suppl. Methods)

### *In vivo* and *in vitro* analysis of (pp)pGpp via HPLC-MS

Nucleotide quantification was performed as described (Gratani *et al*., 2018). Briefly, nucleotides were analyzed using ESI-TOF (micrO-TOF II, Bruker) mass spectrometer connected to an UltiMate 3000 high-performance liquid chromatography. 5 µl of standards or samples were injected onto SEQuant ZIC-pHILIC column (Merck, PEEK 150 × 2.1 mm, 5 µm). MS analysis was performed in negative-ion mode over the mass range from 200 to 1,000 m/z. MS calibration was done by using a sodium formate solution as the tune mix. Nucleotide standards of AMP (346.06 m/z), ATP (505.99 m/z), GTP (521.98 m/z), pGpp (521.98 m/z), ppGpp (601.96 m/z) and pppGpp (681.92 m/z) were diluted 10 times 1:1 from 1 mM until a concentration of 1.9531325 µM and analyzed by HPLC-MS. Extracted ion chromatogram (EIC) spectra of all standards were presented in DataAnalysis (Bruker) and the area under the curve (AUC) of the respective EICs was calculated in GraphPad Prism 5 (baseline was set to 150). The obtained AUC values of the diluted standards were used to generate a calibration curve. For absolute nucleotide quantification, the AUC of the samples was plugged into the AUC values of the calibration curve and the concentration of the respective nucleotides in the samples was determined. Nucleotide identification was verified by matching the retention times and m/z values of detected peaks in the samples to the measured nucleotide standards. To separate pGpp from GTP we used an expectation–maximization (EM). The relative amount of the first chemical component in the mixture is calculated as 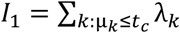. The second component was expressed as *I*_2_ = 1 − *I*_1_..

### H_2_O_2_ killing assay

Strains were grown over night in CDM, diluted in fresh CDM to an OD_600_ = 0.1 and growth followed for 24 hours with different H_2_O_2_ concentrations in a microplate reader (Infinite M200, Tecan). MIC determination was performed according to European Committee on Antimicrobial Susceptibility Testing (EUCAST) guidelines using CDM medium.

### RNA-Seq Analysis

Strains were grown in triplicates to OD_600_ = 0.3, split into treated (ATc, 0.1 µg/ml) and untreated control and grown for 30 min, 37°C. Purified RNA was sent to Vertis Biotechnologie AG Freiburg for RNA Sequencing based on Illumina Next Seq 500 system. RNA was examined by a capillary electrophoresis on a Shimadzu MultiNA microchip followed by rRNA depletion using Ribo-Zero rRNA removel Kit from Illumina. RNA was converted to cDNA by fragmenting RNA samples by ultrasound and ligating an oligonucleotide adapter to the 3’end of the RNA. Using M-MLV reverse transcriptase first strand cDNA was created using 3’ adapter as primer. The 5’Illumina TruSeq sequencing adapter was ligated to the 3’end of the purified (Agencourt AMPure XP kit) cDNA and PCR was performed. Samples were pooled in equimolar amounts and fractionated in a size range of 200-500 bp using a preparative agarose gel and Illumina sequencing was performed using 75 bp reads. RNA-Seq analysis was performed using CLC Genomic Workbench (Qiagen). Reads were trimmed (TrueSeq-Antinsense Primer AGATCGGAAGAGCACACGTCTGAACTCCAGTCA) and mapped to the reference genome of HG001 (NZ_CP018205.1). Differential gene expression was performed comparing RSH-Syn or RelQ versus the (pp)pGpp^0^ mutant. Venn diagrams were performed comparing RSH-Syn vs. control and RelQ vs. control. Genes with at least 3-fold difference and a p-value ≥0.001 were defined as differentially regulated compared to the untreated control.

Data are available with GEO accession GSE145144

## Acknowledgements

We thank Isabell Samp for excellent technical assistance and Ulrich Schoppmeier for support in data analyses. Mutants from the Nebraska library were obtained through the Network on Antimicrobial Resistance in Staphylococcus aureus (NARSA) program. The work was supported by Grants from the Deutsche Forschungsgemeinschaft: TR34/B1 to C.W. and U.M., TRR261/A06 and SFB766/Z01 to C.M., SPP1879 to C. W. and Infrastructural funding from Cluster of Excellence EXC 2124 “Controlling Microbes to Fight Infections”.

