## Supplemental Material for "Inducible expression of (pp)pGpp synthetases in *Staphylococcus aureus* is associated with activation of stress response genes"

### Synthesis of guanosine 3'-*O*-diphosphate 5'-*O*-phosphate pGpp

pGpp was synthesized starting from conveniently protected guanosine and employing both phosphoramidite and phosphotriester methods (Scheme 1).

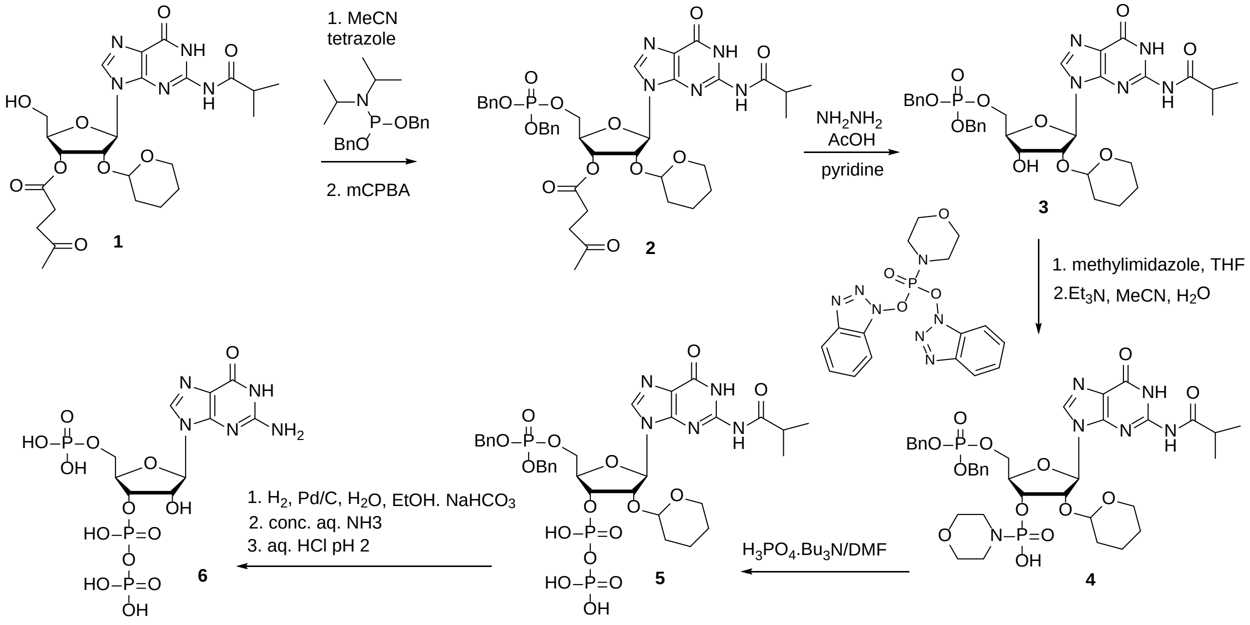


*Scheme 1 Synthesis of pGpp*

### General

Unless stated otherwise, all used solvents were anhydrous. TLC was performed on silica gel pre-coated aluminium plates Silica gel/TLC-cards, UV 254 (Fluka), and compounds were detected by UV light (254 nm)or by spraying with 1% solution of 4-(4-nitrobenzyl)pyridine in ethanol followed by heating and treating with gaseous ammonia (blue color of mono- and diesters of phosphonic acid). Preparative column chromatography was carried out on silica gel (40-60µm; Fluka), and elution was performed at the flow rate of 40 ml/min. Analytical HPLC was performed on reversed phase columns (LUNA Phenomenex, C18) using an LCMS device (Autopurification System, Waters). Preparative RP HPLC was performed on LC5000 Liquid Chromatograph (INGOS-PIKRON, CR) using Luna C18 (2) column (4.6 x 150 mm) at flow rate 1 ml/min by a gradient elution of methanol in 0.1M TEAA pH 7.5 (A = 0.1M TEAA; B = 0.1M TEAA in 50% aqueous methanol; C = methanol). Mass spectra were recorded on LTQ Orbitrap XL (Thermo Fisher Scientific) instrument using ESI method. NMR spectra were measured on Bruker

AVANCE 400 ($^1$H at 400 MHz, $^{13}$C at 100.6 MHz), Bruker AVANCE 500 and Varian UNITY 500 ($^1$H at 500 MHz, $^{13}$C at 125.8 MHz) spectrometers. $D_2O$ (reference (dioxane) = $^1$H 3.75 ppm, $^{13}$C 69.3 ppm. Chemical shifts (in ppm, $\delta$ scale) were referenced to TMS as internal standard; coupling constants (*J*) are given in Hz.

**Dibenzyl (2'-*O*-Tetrahydropyranyl-3'-*O*-levulinoyl-2-*N*-isobutyrylguanosin-5’-yl)phosphate 2 DR-6314**

Tetrazole (11 ml, 0.45M in MeCN) was added to the mixture of 2'-*O*-Tetrahydropyranyl-3'-*O*-levulinoyl-2-*N*-isobutyrylguanosine (prepared according to [1] from 2-*N*-isobutyrylguanosine) (0.54 g, 1 mmol) and dibenzyl *N*,*N*-diisopropylphosphoramidite (0.5 ml, 1.5 mmol) in acetonitrile (10 ml). The reaction mixture was stirred at rt for 1 h under argon atmosphere. 2-Chloroperbenzoic acid (0.34 g, 2 mmol) was added and the reaction mixture was stirred for 20 min. The reaction mixture was concentrated in vacuo and titled product was obtained by chromatography on silica gel using linear gradient of ethanol in chloroform in the form of white foam in 98% yield (0.78 g, 0.98 mmol).

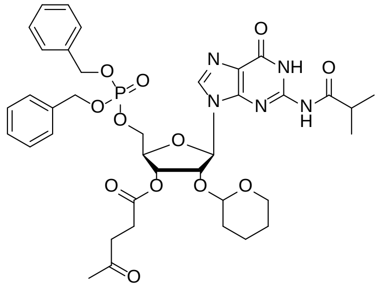

**DR-6314** (#52370)

$^1$H NMR (500.0 MHz, $CDCl_3$): 1.17, 1.21 (2 × d, 2 × 3H, $J_{vic}$ = 6.8, $(C\mathbf{H_3})_2CH$); 1.23-1.31, 1.32-1.45, 1.51-1.61 (3 × m, 3 × 2H, $CH_2$-THP); 2.21 (s, 3H, $CH_3CO$); 2.59 – 2.85 (m, 6H, $(CH_3)_2C\mathbf{H}$, $CH_a\mathbf{H_b}$O-THP, $COC\mathbf{H_2}C\mathbf{H_2}CO$); 2.96 (dt, 1H, $J_{gem}$ = 11.2, $J_{vic}$ = 4.3, $C\mathbf{H_a}H_b$O-THP); 4.32 (t, 2H, $J_{H,P} = J_{5',4'}$ = 4.4, H-5′); 4.36 (m, 1H, H-4′); 4.51 (m, 1H, CHO-THP); 4.94 (dd, 1H, $J_{gem}$ = 12.0, $J_{H,P}$ = 9.4, $CH_a\mathbf{H_b}Ph$); 4.97 (dd, 1H, $J_{gem}$ = 12.0, $J_{H,P}$ = 8.7, $C\mathbf{H_a}H_bPh$); 5.11 (d, 2H, $J_{H,P}$ = 9.2, $CH_2Ph$); 5.24 (dd, 1H, $J_{2',1'}$ = 7.7, $J_{2',3'}$ = 5.4, H-2′); 5.40 (dd, 1H, $J_{3',2'}$ = 5.4, $J_{3',4'}$ = 1.2, H-3′); 5.78

(d, 1H, $J_{1',2'}$ = 7.7, H-1′); 7.08-7.11, 7.19-7.30, 7.36-7.40 (3 × m, 10H, H-*o,m,p*-Ph); 7.67 (s, 1H, H-8); 10.87 (s, 1H, NHCO); 12.17 (s, 1H, NH-1).

$^{13}$C NMR (125.7 MHz, $CDCl_3$): 18.43 ($CH_2$-THP); 18.91, 19.12 ((**C**$H_3$)$_2$CH); 24.73 ($CH_2$-THP); 27.66 (OCO**C**$H_2CH_2COCH_3$); 29.60 ($CH_2$-THP); 29.87 (**C**$H_3$CO); 35.64 (($CH_3$)$_2$**C**H); 37.70 (OCOC$H_2$**C**$H_2COCH_3$); 61.14 ($CH_2$O-THP); 66.82 (d, $J_{C,P}$ = 6.3, $CH_2$-5′); 69.39, 69.94 (2 × d, $J_{C,P}$ = 5.7, $CH_2$Ph); 71.16 (CH-3′); 74.26 (CH-2′); 81.12 (d, $J_{C,P}$ = 9.2, CH-4′); 88.73 (CH-1′); 97.90 (CHO-THP); 122.81 (C-5); 127.18, 128.18, 128.55, 128.74, 128.97 (CH-*o,m,p*-Ph); 134.90, 135.16 (2 × d, $J_{C,P}$ = 6.0, C-*i*-Ph); 139.53 (CH-8); 147.93 (C-2); 148.15 (C-4); 155.61 (C-6); 171.87 (COO); 179.99 (NOC); 206.21 (**C**O$CH_3$).

$^{31}$P{$^1$H} NMR (202.3 MHz, $CDCl_3$): -1.43.

**HR-ESI** $C_{38}H_{47}O_{12}N_5P$ $(M+H)^+$ calcd. 796.29533, found 796.29562.

**(2'-*O*-Tetrahydropyranyl-2-*N*-isobutyrylguanosine 5’-(dibenzyl phosphate) 3 DR-6315**

Compound 2 was dissolved in the mixture of pyridine (10 ml), acetic acid (5 ml) and hydrazine hydrate (0.7 ml) and stirred at rt for 30 min. The reaction mixture was diluted with chloroform (50 ml) and washed with sat. soln. of $NaHCO_3$. The organic phase was dried with $Na_2SO_4$, filtered and concentrated in vacuo. Titled compound was obtained by chromatography on silica gel using linear gradient of ethanol in chloroform in the form of white foam in 78% yield (0.53 g, 0.76 mmol).

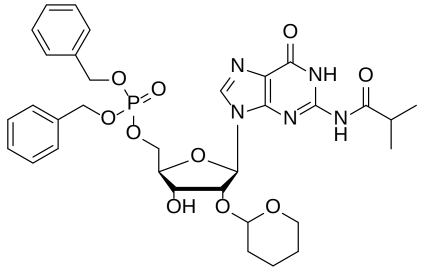


**DR-6315** (#52371)

$^1$H NMR (500.0 MHz, $CDCl_3$): 1.17, 1.22 (2 × d, 2 × 3H, $J_{vic}$ = 6.8, (C**H**$_3$)$_2$CH); 1.33-1.48, 1.50-1.58, 1.64-1.74 (3 × m, 3 × 2H, $CH_2$-THP); 2.74 (quin, 1H, $J_{vic}$ = 6.8, ($CH_3$)$_2$C**H**); 2.94 (bs, 1H, OH-3′); 3.04 (dt, 1H, $J_{gem}$ = 11.2, $J_{vic}$ = 6.6, 3.9, $CH_a$**H**$_b$O-THP); 3.26 (dt, 1H, $J_{gem}$ = 11.2, $J_{vic}$ =

7.3, 3.7, C**H$_a$**H$_b$O-THP); 4.22-4.38 (m, 2H, H-4′,5′b); 4.37 (m, 1H, H-5′a); 4.49-4.52 (m, 1H, H-3′, CHO-THP); 4.95 (dd, 1H, $J_{gem}$ = 12.0, $J_{H,P}$ = 9.2, CH$_a$**H$_b$**Ph); 4.98 (dd, 1H, $J_{gem}$ = 12.0, $J_{H,P}$ = 8.6, C**H$_a$**H$_b$Ph); 5.02 (dd, 1H, $J_{2',1'}$ = 6.3, $J_{2',3'}$ = 5.5, H-2′); 5.12 (dd, 1H, $J_{gem}$ = 11.7, $J_{H,P}$ = 9.0, CH$_a$**H$_b$**Ph); 5.16 (dd, 1H, $J_{gem}$ = 11.7, $J_{H,P}$ = 9.0, C**H$_a$**H$_b$Ph); 5.88 (d, 1H, $J_{1',2'}$ = 6.3, H-1′); 7.11-7.14, 7.22-7.31, 7.35-7.44 (3 × m, 10H, H-*o,m,p*-Ph); 7.65 (s, 1H, H-8); 10.88 (s, 1H, NHCO); 12.17 (s, 1H, NH-1).

$^{13}$C NMR (125.7 MHz, CDCl$_3$): 18.90, 19.15 ((**C**H$_3$)$_2$CH); 19.54, 24.65, 30.25 (CH$_2$-THP); 35.64 ((CH$_3$)$_2$**C**H); 63.05 (CH$_2$O-THP); 67.02 (d, $J_{C,P}$ = 6.5, CH$_2$-5′); 69.39, 69.87 (2 × d, $J_{C,P}$ = 5.7, CH$_2$Ph); 70.15 (CH-3′); 77.75 (CH-2′); 82.82 (d, $J_{C,P}$ = 8.5, CH-4′); 89.11 (CH-1′); 99.90 (CHO-THP); 122.63 (C-5); 127.24, 128.16, 128.56, 128.71, 128.73, 128.92 (CH-*o,m,p*-Ph); 134.94, 135.30 (2 × d, $J_{C,P}$ = 6.2, C-*i*-Ph); 139.69 (CH-8); 147.85 (C-2); 148.10 (C-4); 155.65 (C-6); 179.98 (NOC).

$^{31}$P{$^1$H} NMR (202.3 MHz, CDCl$_3$): -1.35.

**HR-ESI** $C_{33}H_{41}O_{10}N_5P$ $(M+H)^+$ calcd. 698.25856, found 698.25876.

**2'-*O*-Tetrahydropyranyl-2-*N*-isobutyrylguanosine 3'-*O*-(phosphoromorpholidate) 5’-*O*-( dibenzyl phosphate) 4**

A solution of *O,O*-bis[l-benzotriazolyl]phosphoromorpholidate in THF (0.2 M; 7.5 ml) and *N*-methylimidazole (0.15 ml; 1.9 mmol) was added to the solution of **3** in THF (10 ml). The reaction mixture was concentrated under the exclusion of moisture until the residual volume was ca. 5 ml and left for 2 h at room temperature. LCMS indicated complete conversion of the starting material. The reaction mixture was diluted at 0 °C with methylene chloride (40 ml) and washed at 0 °C with an aqueous solution of triethylammonium bicarbonate (0.5 M; 3 x 10 ml), water (2 x 15 ml), an aqueous solution of potassium dihydrogen phosphate (0.5 M; 2 x 15 ml, pH 6) and water (2 x 15 ml). The organic layer was dried with $Na_2SO_4$ (5 g) and concentrated to afford a colourless oil which was triturated with petroleum-ether (40-60 °C; 3 x 50 ml). Obtained intermediate was dissolved in acetonitrile (5 ml). Triethylamine (2.5 ml) and water (1.25 ml) were added. After standing for 4 h at room temperature, LCMS indicated complete conversion of the starting compound into baseline-material. Toluene (50 ml) was added and the resulting solution was concentrated under diminished pressure at 25 °C to afford pure 2'-*O*-Tetrahydropyranyl-2-*N*-isobutyrylguanosine 3'-*O*-(phosphoromorpholidate) 5'-*O*-(dinebzyl phosphate) as a white glass that was characterized with LCMS and used without further characterization in the next step.

**2'-*O*-Tetrahydropyranyl-2-*N*-isobutyrylguanosine 3'-*O*-diphosphate 5'-*O*-(dibenzyl phosphate) 5**

A solution of mono-(tri-n-butylammonium) phosphate in dimethylformamide (0.5 M; 6 ml) was added to the solution of **4** in DMF (5 ml). The reaction mixture was rendered anhydrous by repeated co-evaporation with toluene (3 x 25ml). After 18 h at 50 °C, the reaction mixture was neutralized with tri-n-butylamine until the pH had become 8.5. The resulting solution was concentrated to a small volume (3 ml) and purified with HPLC on reversed phase using linear gradient of methanol in 0.1M aqueous TEAB. The product was identified by means of LCMS and used without further characterization in the next step.

**guanosine 3'-*O*-diphosphate 5'-*O*-phosphate 6**

Palladium on charcoal (20 mg) was added to solution of compound 5 in water. The mixture was hydrogenated at 100 kPa of $H_2$ overnight. The reaction mixture was filtrated over a pad of cellite, concentrated in high vacuum at 25 °C. Concentrated ammonia (14.8 M; 50 ml) was added and the resulting clear solution was left for 24 h at rt. LCMS indicated complete removal of the iBu group. The solvents were evaporated in high vacuum at 25 °C, and the residue and dissolved in hydrochloric acid (0.01 N; 2 ml). The pH of the solution was adjusted to 2.00 with hydrochloric acid (0.1 N). The clear solution was left for 1 h at 20 °C, and then carefully neutralized with dilute ammonia (0.5 M) until the pH of the solution had become 8.00. Excess ammonia was evaporated and the residue (1 ml) was applied on the column of reversed phase. The final product 6 was obtained by preparative HPLC on reversed phase using linear gradient of methanol in 0.1M aqueous TEAB and converted to ammonium salt by passing through a small column of Dowex 50 in $NH_4^+$ cycle. After lyophilisation from water 150 mg (0.25 mmol, 33% over five steps) of pGpp was obtained as a white amorphous solid.

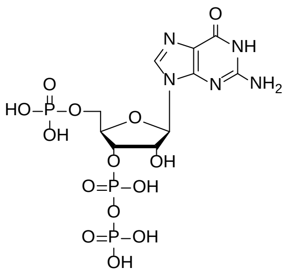

**pGpp DR-6050** (#50231)

**$^{1}$H NMR** (500.0 MHz, $D_2O$, ref(dioxane) = 3.75 ppm) δ 4.09 – 4.17 (m, 2H, H-5′), 4.56 (dt, $J_{4′,5′}$ = 5.2, 2.9, $J_{4′,3′}$ = 2.9, 1H, H-4′), 4.86 (ddd, $J_{2′,1′}$ = 6.4, $J_{2′,3′}$ = 5.2, $J_{H,P}$ = 1.3, 1H, H-2′), 4.95 (ddd, $J_{H,P}$ = 8.6, $J_{3′,2′}$ = 5.2, $J_{3′,4′}$ = 2.9 Hz, 1H, H-3′), 5.99 (d, $J_{1′,2′}$ = 6.4 Hz, 1H, H-1′), 8.11 (s, 1H, H-8).

**$^{13}$C NMR** (125.7 MHz, $D_2O$, ref(dioxane) = 69.3 ppm) δ 67.17 (d, $J_{C,P}$ = 4.9 Hz, $CH_2$-5′), 75.83 (d, $J_{C,P}$ = 4.8 Hz, CH-2′), 77.56 (d, $J_{C,P}$ = 5.4 Hz, CH-3′), 85.97 (dd, $J_{C,P}$ = 8.7, 3.7 Hz, CH-4′), 89.39 (CH-1′), 119.02 (C-5), 140.48 (CH-8), 154.64 (C-4), 156.73 (C-2), 161.79 (C-6).

**$^{31}$P NMR** (202.3 MHz, $D_2O$) δ -10.44 (dd, $J_{P,P}$ = 20.7, $J_{P,H}$ = 8.6 Hz, Pα-3′), -8.71 (d, $J$ = 20.7 Hz, Pβ-3′), 1.45 (s, P-5′).

**HR-ESI** $C_{10}H_{15}O_{14}N_5P_3$ $(M\text{-}H)^-$ calcd. 521.98338, found 521.98312.

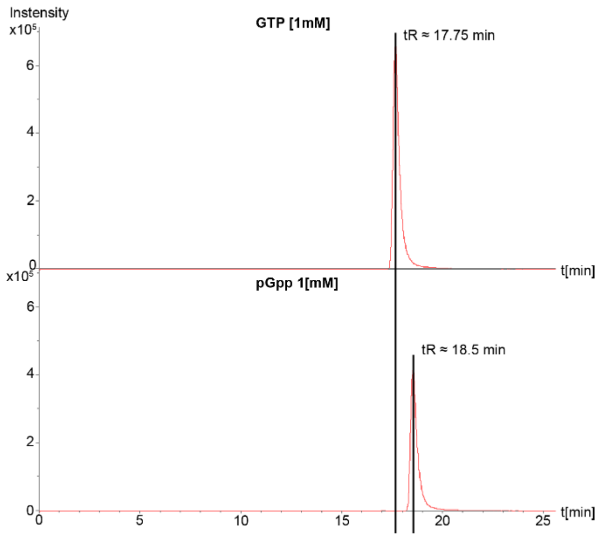


Elutionsprofile of GTP versus pGpp analyzed using ESI-TOF (micrO-TOF II, Bruker) mass spectrometer connected to an UltiMate 3000 high-performance liquid chromatography analyzed using ESI-TOF (micrO-TOF II, Bruker) mass spectrometer connected to an UltiMate 3000 high-performance liquid chromatography.

1. Schattenkerk, C.; Wreesmann, C. T. J.; van der Marel, G. A.; van Boom, J. H., Synthesis of riboguanosine pentaphosphate ppprGpp (Magic Spot II) via a phosphotriester approach. *Nucleic Acids Research* **1985,** *13* (10), 3635-3649.

**Table S3: Strains and plasmids**

| Strains | Description | Source/ Reference |
|---|---|---|
| ***E.Coli*** | | |
| BL21 | fhuA2 [lon] ompT gal (λ DE3) [dcm] ΔhsdS λ DE3= λ sBamHlo ΔEcoRI-B int:: (lacl::PlacUV5::T7 gene1) i21 Δnin5, competent for protein expression | NEB |
| ***S.aureus*** | | |
| RN4220 | Restriction deficient derivate of 8325-4, $rK^{-}mK^{+}$ | (Kreiswirth et al., 1983) |
| Ne1193 | Tnbursa::*sarA erm* | NARSA |
| NE99 | Tnbursa::*fur erm* | NARSA |
| NE665 | Tnbursa::*perR erm* | NARSA |
| HG001 | RN1 derivate, rsbU repaired, tcaR | (Pohl et al., 2009)<br>(Herbert et al., 2010) |
| HG001-86 | Mutation in the synthetase domain of *rel* | (Geiger et al., 2010) |
| HG001-229-230 | Mutation in the synthetase domain of *relP* and *relQ* ($\Delta relP_{syn}$ $\Delta relQ_{syn}$) | (Geiger et al., 2014) |
| HG001 229-230-263 | Mutation in the synthetase domain of *relP* ,*relQ* and complete deletion of *rel* ($\Delta relP_{syn}$ $\Delta relQ_{syn}$ Δrel) | (Geiger et al., 2014) |
| HG001 *fur* | Tnbursa::*fur erm* | This work |
| HG001 229-230-263 *fur* | Mutation in the synthetase domain of *relP* ,*relQ* and complete deletion of *rel* ($\Delta relP_{syn}$ $\Delta relQ_{syn}$ Δrel) Tnbursa::*fur erm* | This work |
| HG001 *perR* | Tnbursa::*perR erm* | This work |

| | | |
|---|---|---|
| HG001 229-230-263 *perR* | Mutation in the synthetase domain of *relP* ,*relQ* and complete deletion of *rel* ($\Delta relP_{syn}$ $\Delta relQ_{syn}$ $\Delta rel$) Tnbursa::*perR erm* | This work |
| HG001 *psmα psmβ* | *psmα1-4::tetM, psmβ1-2::ermC* | (Geiger et al., 2012) |
| HG001 229-230-263 *psmα psmβ* | Mutation in the synthetase domain of *relP* ,*relQ* and complete deletion of *rel* ($\Delta relP_{syn}$ $\Delta relQ_{syn}$ $\Delta rel$) *psmα1-4::tetM, psmβ1-2::ermC* | This work |
| USA300 JE2 | USA300 derivative, cured of all plasmids | NARSA |
| USA300 JE2 229-230-263 | Mutation in the synthetase domain of *relP* ,*relQ* and complete deletion of *rel* ($\Delta relP_{syn}$ $\Delta relQ_{syn}$ $\Delta rel$) | This work |
| **Plasmids** | **Description** | **Source/ Reference** |
| pET15b | Protein expression vector, ampicillin resistance | Novagen |
| pKOR1 | ATc-inducible mutagenisis vector, chloramphenicol resistance | (Bae & Schneewind, 2006) |
| pCG248 | anhydrotetracyclin (ATc) inducible vector, chloramphenicol resistance | (Helle et al., 2011)<br>(Schroder, Goerke, & Wolz, 2013) |
| pCG258 | relP cloned into pCG248 | (Geiger et al., 2014) |
| pCG259 | relQ cloned into pCG248 | (Geiger et al., 2014) |
| pCG327 | N-terminal domain of Rel with hydrolase mutated | (Gratani et al., 2018) |
| pCG229 | pKOR1 with integrated, mutated *relP* | (Geiger et al., 2014) |

| | | |
|---|---|---|
| pCG230 | pKOR1 with integrated, mutated *relQ* | (Geiger et al., 2014) |
| pCG263 | pKOR1 with integrated, mutated *rel* | (Geiger et al., 2014) |
| pCG551 | N-terminal domain of Rel with hydrolase mutated cloned into pET15b | (Gratani et al., 2018) |
| pCG121 | relP cloned into pET15b | (Geiger et al., 2014) |
| pCG122 | relQ cloned into pET15b | (Geiger et al., 2014) |

**Table S4: Oligonucleotides**

| Purpose and Description | Template | Name | Sequence |
|---|---|---|---|
| verification of *relP* synthase mutant | USA300-229-230-263 | relPDIG-for<br>relPDIG-rev | GTCGCACATTCTTTCAGT<br>CGTTATTAGGTTTCGTAGAGTT |
| verification of *relQ* synthase mutant | USA300-229-230-263 | relQDIGfor2<br>relQDIGrev2 | TTCGTAACACTAAAGAAAGTGG<br>GCGTGTAATATTTTTGAGCT |
| verification of *rsh* mutant | USA300-229-230-263 | rel431for<br>relLC4rev | GCGTGGCTTTATCATTGG<br>ACTTCAACCATCATTCGG |
| Verification of *perR* mutant | HG001 *perR*<br>HG001 229-230-263 *perR* | perR-for<br>TnUpstream | TGAACTAGAAGAATCAATTGCATCA<br>CTCGATTCTATTAACAAGGG |
| Verification of *fur* mutant | HG001 *fur*<br>HG001 229-230-263 *fur* | furtnfor<br>TnBuster | GCACGTTTCACACACACCAT<br>GCTTTTTCTAAATGTTTTTTAAGTAAATCAAGTAC |
| Verification of *sarA* mutant | HG001 *sarA*<br>HG001 229-230-263 *sarA* | sarAtnfor<br>TnUpstream | GTTGTTTGCTTCAGTGATTCGT<br>CTCGATTCTATTAACAAGGG |
| Verification of *psmα/β* mutant | HG001 229-230-263 *psm* | | (Geiger et al., 2012) |
| Creation of dig-labeled probe *ftnA* | WT HG001 | ftnADig-for<br>ftnADig-rev | GAGTACTTTGCAGCACACGC<br>CATTGCTGTCATCGCCGATAC |
| Creation of dig-labeled probe *dps* | WT HG001 | mrgADig-for<br>mrgADig-rev | GCTACACAATTTCCACTGGT<br>CATACCTATAAACATATCTTC |
| Creation of dig-labeled probe *psm*/*agr* | | | (Geiger et al., 2012) |
